# Host-parasite coevolution can promote the evolution of seed banking as a bet-hedging strategy

**DOI:** 10.1101/273003

**Authors:** Mélissa Verin, Aurélien Tellier

**Author notes:** Contributions: MV and AT designed the study, performed the analytical computations, and wrote the manuscript. MV performed the numerical simulations and analysed the results.

## Abstract

Seed (egg) banking is a common bet-hedging strategy maximizing the fitness of organisms facing environmental unpredictability by the delayed emergence of offspring. Yet, this condition often requires fast and drastic stochastic shifts between good and bad years. We hypothesize that the host seed banking strategy can evolve in response to coevolution with parasites because the coevolutionary cycles promote a gradually changing environment over longer times than seed persistence. We study the evolution of host germination fraction as a quantitative trait using both pairwise competition and multiple mutant competition methods, while the germination locus can be genetically linked or unlinked with the host locus under coevolution. In a gene-for-gene model of coevolution, hosts evolve a seed bank strategy under unstable coevolutionary cycles promoted by moderate to high costs of resistance or strong disease severity. Moreover, when assuming genetic linkage between coevolving and germination loci, the resistant genotype always evolves seed banking in contrast to susceptible hosts. Under a matching-allele interaction, both hosts’ genotypes exhibit the same seed banking strategy irrespective of the genetic linkage between loci. We suggest host-parasite coevolution as an additional hypothesis for the evolution of seed banking as a temporal bet-hedging strategy.

## Introduction

Seed (egg) banking consists in the variation in the timing of emergence of viable seeds or eggs from a single clutch that are stored in the soil, river or lake sediments (Evans & Dennehy 2005). This life history strategy is common to many plants (Venable 1989; Philippi 1993; Clauss & Venable 2000) insects (Hanski 1988; Gourbière & Menu 2009) and crustaceans (Moustakas & Evans 2013). Only a fraction of seeds germinates each year, decreasing the population growth rate when environmental conditions are favourable while avoiding extinction when conditions are drastic. Seed banking is expected to evolve in stochastic and unpredictable environments as a temporal bet-hedging strategy (Boer 1968; Slatkin 1974; Seger & Brockmann, 1987; Philippi & Seger, 1989), spreading the risk of reproductive failure through time dispersion. This strategy is also advantageous to mitigate the negative impact of sibling competition, overcrowding and inbreeding (Westoby 1981; Kobayashi & Yamamura 2000).

In the context of bet-hedging theory, classical models investigate the evolution of the optimal germination fraction (which we denote here *b*_*0*_). This fraction defines the proportion of seeds produced by a plant that germinates each season, as opposed to the proportion of seeds entering the bank (1−*b*_*0*_) (Cohen 1966; Bulmer 1984; Ellner 1985; Valleriani 2005). According to these studies, the optimal germination fraction is expected to be low (high fraction of seeds entering the seed bank) for highly variable environments, such as variation in water availability or other abiotic factors, but also disturbance of habitats. The models cited above consider drastic environmental contexts, in particular annual plants growing in deserts, where the environment is represented as a fast succession of good and bad conditions. The reproduction and/or survival of plants is optimal during good seasons and strongly reduced (as far as null) during bad seasons. However, seed banking plants are widespread and observed in temperate climate where environmental changes are more gradual.

Another interesting life history characteristic generating variable environments over time is the interaction between hosts and their parasites or parasitoids. It has been well documented that such interactions promote coevolutionary dynamics resulting in fluctuating selection, fixation of alleles or stable polymorphism in both species (*e.g.* (Leonard 1977; Gandon *et al.* 1996; Tellier & Brown 2007b; Ashby & Boots 2017). The dynamics are driven by negative indirect frequency-dependent selection (niFDS) – rare alleles exhibiting fitness advantages as selection in the host population depends on allele frequencies in the parasite population, and *vice versa* (Clarke 1964; Tellier & Brown 2007b). We investigate in this article the context of unstable coevolutionary dynamics, with cycles increasing over time in their amplitude and period (Leonard 1977; Tellier & Brown 2007b, 2009). These unstable cycles are therefore unpredictable for hosts and parasites in time, and their characteristics are further dependent on the genetic interaction between host and parasite. In the theoretical literature, the genetic determinism of host-parasite interactions has traditionally been modelled in two ways (but see van Baalen 1998; Gandon *et al.* 2002; Boots *et al.* 2014; Ashby & Boots 2017). Either host and parasite genotypes are specific to one another and must match for the infection to be successful (Matching allele, MA, (*e.g.* Gandon *et al.* 1996; Agrawal & Lively 2002; Dybdahl *et al.* 2014), or genotypes vary in their degree of specialization, from specialists to generalists (Gene for gene, GFG) resulting in more complex coevolutionary dynamics (*e.g.* Flor 1971; Leonard 1977; Frank 1992; Agrawal & Lively 2002; Dybdahl *et al.* 2014). We hypothesize here that unstable coevolutionary dynamics generate the unpredictable variation necessary for evolving seed banking as a bet-hedging strategy in hosts.

This specific co-evolutionary mechanism for the evolution of temporal bet-hedging has not yet been explored, while it was suggested in a review article by Hanski (1988). He indeed questioned whether cyclic or chaotic host/parasitoid and predator/prey dynamics can promote the evolution of extra long diapause of insects (*i.e.* equivalent to seed banking). Note that the reverse mechanism has been well studied, namely that prolonged diapause (or seed banking) can stabilise host-parasitoid and host-parasite dynamics (Tellier & Brown 2009, Ringel *et al.* 1998). Though, the stabilizing property relies on the germination of seeds to be geometrically distributed and the time in the bank to be unbounded, such that an egg or a seed can remain infinitely dormant with a constant germination probability (Corley *et al.* 2004). Few theoretical studies assume a bounded seed bank (of more than two years) (Templeton & Levin 1979; Valleriani & Tielbörger 2006), however, in biology, seed banks are characterised by their persistence and accordingly classified in three categories (Fenner & Thompson 2005). Transient seed banks are formed by seeds surviving less than a year in the soil before their decay. Short term and long term persistent banks are, respectively, formed by seeds surviving less and more than five years. Three characteristics compose a seed banking strategy: the germination fraction (b_0_), the shape of the germination function (often assumed to be geometric) and whether the life spans of a seed is finite (bounded to a maximum value).

We propose a general model to test Hanski’s intuition in the context of host-parasite coevolution and aim to demonstrate an additional mechanism generating the evolution of seed banking as temporal bet-hedging, namely the evolution of a strategy that maximises the geometric fitness of hosts under parasite pressure. Our model assumes an infinite host and parasite population, each composed of two types (resistant/susceptible and infective/non-infective), and includes a short or long term persistent bank for the host. We firstly assess that the geometric mean fitness criteria, determining which allele goes to fixation in an infinite population model (Templeton and Levin 1979, Starrfelt & Kokko 2012) holds for models assuming the competition of two alleles (*i.e.* two hosts types). Then using both pairwise competition and multiple competition simulations of seed banking strategies, we show that distinct germination fractions emerge depending on the genetic linkage between the germination and coevolutionary loci and the considered model of genetic interaction (MA or GFG). We expose a complex interaction between the properties of the coevolutionary cycles (period and amplitude of the fluctuations) and the characteristics of the seed banking strategies. Finally, this allows us to extend previous classical results in finite size models, and to study temporal bet-hedging as an eco-evolutionary adaptation to slow but sustained biotic environmental fluctuations.

## Materials and methods

### Model description

We assume an infinite population size model describing a GFG interaction between hosts and parasites (*e.g.* Tellier & Brown 2009). For simplicity, both organisms are haploid as models with diploids generate similar behaviour and dynamics (Ye *et al.* 2003). One genetic locus with two alleles describes the GFG interaction, with the host exhibiting a resistant (*R*) or a susceptible (*r*) allele, and the parasite an infective (*INF*) or a non-infective (*ninf*) allele. At generation *g* the resistant and susceptible hosts have frequencies *r*_*g*_ and *R*_*g*_ respectively, and the infective and non-infective parasites, *a*_*g*_ and *A*_*g*_ respectively. Host frequencies are scaled by the mean fitness of the host population 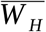, and parasite frequencies by the mean fitness of the parasite population 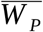. Carrying resistance or infectivity alleles comes with the costs *C*_*H*_ and *C*_P_. When infected, host fitness is reduced by *s*, the so-called disease severity. Non-infective parasites cannot infect resistant hosts. We assume that each host is exposed to parasites at each generation (Leonard 1977, Tellier and Brown 2007). The following difference equations describe changes in allele frequencies with a seed bank for the host.
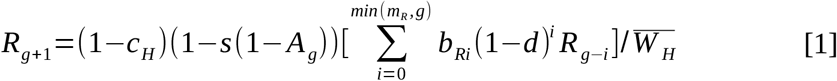

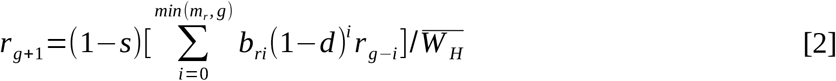

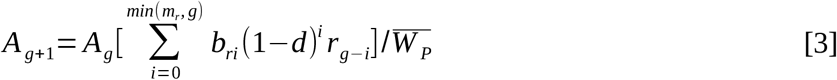

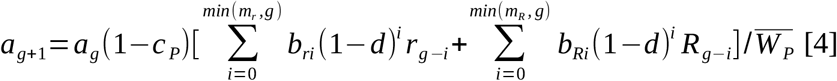

The seed bank is modelled based on a forward in time adaptation of the model from (Kaj *et al.* 2001). In general terms, a distinct quantitative locus determines the rate *b*_*i*_ at which seeds produced *i* generations ago germinate at a given generation *g*. The maximum amount of time seeds can remain in the bank is fixed to *m*. A key parameter of the seed bank is the fraction *b*_*0*_ of newly produced seeds that germinate in the next generation, if b_0_ =1 all seeds are non-dormant and the bank is empty. A germination function describes the relative contribution of non-dormant and dormant seeds to the above ground population, *b*_*0*_ versus *b*_*i*≠*0*_. We assume a geometric memoryless function of germination, *b*_*i*_ = *b*_*0*_(1−*b*_*0*_)^(*i*−1)^ (Fig. S1). In other words, the time spent in the bank does not affect the germination rate of the seed *per se*, but older seeds contribute less to the above ground population. Note that by definition, 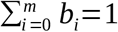 and the germination function is truncated. The germination function is applied to the resistant and susceptible hosts which thus have the respective rate and seed bank persistence *b*_*Ri*_ and *m*_*R*_, and *b*_*ri*_ and *m*_*r*_. Seed banking comes with a cost *d*, the rate at which seed dies per generation. We choose an intuitive geometric seed death rate function, so the probability for a seed to be viable and able to germinate after *i* generations is (1−*d*)_*i*_. The rate of seed death, *d*, is identical for *R* and *r* alleles.

Our model is analogous to a modifier model (e.g. Blanquart & Gandon 2011), in which the host genome is composed of two distinct loci, either linked by being physically close on a single chromosome, or unlinked by being far apart on the same chromosome or being on different chromosomes. The modifier locus is quantitative and controls the germination strategy of the host, and the second locus is under fluctuating selection due to coevolution. We do not assume a seed bank for the parasite, its genome being reduced to the single corresponding locus under fluctuating selection.

The questions we address are 1) whether the coevolutionary dynamics select for the evolution of a seed banking strategy, 2) whether an optimal germination rate exists in that case, and 3) if the optimal strategy differs for resistant and susceptible hosts. The evolution of the seed banking strategy is the result of a change in either the germination fractions *b*_*R0*_ and/or *b*_*r0*_, or the length of the seed bank *m*_*R*_ and/or *m*_*r*_. Both changes lead to a new distribution of the germination function *b*_*i*_ (based on the geometric function) and modify the characteristics of the coevolutionary cycles. We focus here on the evolution of *b*_*0*_, as we later explain (and had confirmed in preliminary simulations, data not shown) that the seed bank persistence *m* would raise to extremely high values, that may not be physiologically relevant for a seed. Due to the feedback between the evolving germination rate *b*_*0*_ and the coevolutionary dynamics, we cannot approximate analytically the period and amplitude of the unstable coevolutionary cycles and therefore we use two methods of simulations.

### Evolution of seed banking strategy: pairwise competition

Our first method investigates the fate of one mutant, denoted *b*_*0*_*, in a resident population with strategy *b*_*0*_. We perform pairwise competition between the mutant and the resident drawing an analogy to pairwise invasion plots used in adaptive dynamics approaches. The mutant *b*_*0*_* is introduced with frequency 0.01 in the population after a burn in phase of 20,000 generations. The burn in phase allows to replenish the seed bank and run several coevolutionary cycles. Susceptible and resistant hosts have here the same mutant (*b*_*0*_*=*b*_*r0*_*=*b*_*R0*_*), and resident (*b*_*0*_=*b*_*r0*_=*b*_*R0*_) strategies. Note however, that due to the coevolutionary dynamics a given mutant may invade only one or both host types. We perform competitions between the whole continuum of seed banking strategies, with *b*_*0*_ ranging from 0.01 to 1, and simulate the dynamics for 1,500,000 generations. This time is chosen to be long enough for fixation or loss of the mutant strategy and is much longer than that used for studying only coevolutionary cycles (Tellier & Brown 2009). We consider the mutant invasion to be successful when its mean frequency is higher than 0.9 over the last 10,000 generations. The mutant and resident strategies are denoted as coexisting when both are present with mean frequency higher than 0.1 and lower than 0.9 in this same time interval.

### Evolution of seed banking strategy: multiple mutant competition

As an alternative method to investigate the process of successive mutation events, we perform simulations of competition between multiple mutants (see for example Boots *et al.* 2014). After a burn in phase of 20,000 generations, a number *N* of mutants with different strategies (different values of *b*_*0*_*) are introduced in a resident population (with initial value *b*_*0*_). We use here *N*=5 (in Boots *et al.* 2014, *N*=1). Each mutant has a different seed banking strategy, *b*_*0*_*, which is sampled in a Normal distribution with mean the resident value *b*_*0*_ and a small standard deviation σ=0.05. The *N* mutants are introduced with equal frequencies summing to 0.01 (so in effect an introduction frequency of 0.01/*N* for each mutant). The resident and the mutants compete during a fixed number of generation *T*=1,000, and we then compute the geometric mean fitness over *T* for each strategy. The strategy with the smallest fitness amongst the *N*+*1* genotypes present (*N b_*0*_*\* mutants and the *b*_*0*_ resident) is removed from the population, and a new mutant with frequency 0.01/*N* is introduced. We make the assumption that hosts with the highest fitness have higher chances to produce mutants, and that the mutational step is small. Thus the new mutant strategy *b*_*0*_* to be introduced is sampled in a Normal distribution with a mean equal to the population average strategy 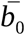 and a small standard deviation σ=0.05. The average strategy 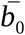 is defined as the sum of the remaining strategies (*N* values of *b*_*0*_) times their geometric mean of frequency in the population.

Two cases are investigated, (i) independence and (ii) non-independence of germination and resistance loci (*e.g.* linkage equilibrium or disequilibrium). In other words, either resistant and susceptible types have the same germination strategy which is then the strategy of the population (*b*_*0*_=*b*_*r0*_=*b*_*R0*_), or these alleles can evolve each their own strategy (*b*_*r0*_≠*b*_*R0*_). The method described above corresponds to the first case (linkage equilibrium). In the case of linked loci, we amend the above simulation protocol by removing and then adding at a given time point one resistant and one susceptible mutant. The mutant strategies are sampled around the average strategy of each type 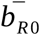 and 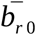, respectively.

We investigate the evolution of seed banking under coevolution using the two simulation methods described above. We contrast the evolution of seed banking for different parameter combinations: high and low costs *C*_*H*_ and *C*_*P*_ (for values of 0.05, 0.1, 0.2 and 0.4), low to high disease severity (*s*=0.1, 0.3, 0.6, and 0.9) and under short term or long term seed bank persistence *m* (5 or 15 years). In addition, we test two hypotheses regarding the genetic linkage between the host locus for coevolution and the locus for the seed banking trait (determining the germination rate *b*_*i*_). For each set of parameters, we perform 50 repetitions, and record over 2×10^6^ generations the population mean strategy 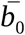, or the resistant and susceptible mean strategies 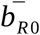 and 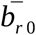 to account for the variability of the mutation sampling procedure. The code for simulation is available in the SI files.

## Results

### Preliminary analysis: bet-hedging in infinite population models

In infinite population models, the geometric mean fitness of a genotype determines which allele goes to fixation (Templeton and Levin 1979, Starrfelt & Kokko 2012). This criterion holds for models of competition between two alleles. Indeed an allele 1 exhibiting randomly varying fitness over time outcompetes an allele 2 with constant fitness if the geometric mean fitness of allele 1 is higher than the relative fitness of allele 2 (SI text S1). If both alleles show stochastic fitnesses over time but exhibit different seed banking strategies (SI text S2), we show that the allele reaching fixation has the strategy maximizing its geometric mean fitness. Our key assumption states that in a deterministic seed bank, the relative proportion of seeds from a given past generation depends on the population fitness at that generation. These results are partially described in the literature (see the more rigorous work by Templeton and Levin 1979). We nevertheless recapitulate them here to introduce our system of notations and extend the previous results in the finite population models by Cohen (1966) and Lewontin & Cohen (1969) to an infinite population.

Templeton and Levin (1979) further highlight the consequences of cyclically varying environments (*i.e.* deterministic environmental cycles) on the fitness of two competing alleles. They identify a key parameter: the ratio between the seed bank persistence and the period of the environmental cycles. They investigate rather short cycling periods (*e.g.* 3 years), however environmental fluctuations in temperate climate may likely vary over longer periods of time. Assuming a cyclic environment and a single host population with a seed banking strategy, we investigate the influence of the seed bank persistence *m* on the optimal germination fraction *b*_*0*_ as a function of the period and amplitude of environmental variations (SI text S3). As expected (Templeton and Levin 1979), for seed bank persistence exceeding the period of environmental variation the host population shows a high investment in the seed bank seen as a higher geometric mean fitness and a low optimizing germination fraction (Fig. S2A). However, when assuming that seed persistence is smaller than the period of drastic environmental variation, we find that low germination fraction can still be advantageous (Fig S2B). With cycles of reduced amplitude, the fitness differences between the various persistence values are drastically reduced, and low germination fractions are only likely to evolve when *m* is equal to the environmental period (Fig. S2C). In the special case of a short term persistent bank (*m*=5), we observe high optimal germination fractions corresponding to weak investment up to no (*b_0_* = 1) investment in the bank, irrespective of the amplitude of the environmental variations (Fig S2B, and C).

In contrast to these cases with regular cycles, optimality theory can not be applied in our model of host-parasite coevolution, since the coevolutionary cycles show 1) increasing period and amplitude over time, and 2) these are continuously shaped by the evolution of the seed banking strategies.

### Dynamical system behavior and seed bank

The system of equations (1) of GFG coevolution with seed banking has several equilibrium points defined for any germination function, which occur as 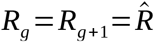 and 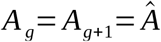. Four so-called trivial points exist as the four monomorphic outcomes (0,0), (1,0), (0,1) and (1,1) for the frequency of *R* and *ninf* (Leonard 1977). There is also one non-trivial equilibrium at which all four host and parasite alleles are maintained: 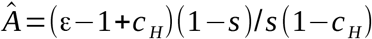 and 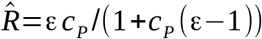. The ratio of the relative germination functions for *R* and *r* alleles including seed mortality is 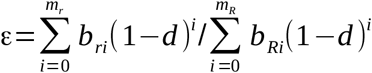, which is equal to one if the resistant and susceptible alleles have the same germination function and seed bank persistence. The internal equilibrium point exists if the parameters fulfil two conditions. First, *C*_*H*_ >1 − ε meaning that the susceptible allele has a germination rate (including seed mortality) equal or slightly smaller than that of the resistant allele (as *C*_*H*_ is assumed to be realistically small, Leonard 1977, Tellier and Brown 2007). Second, *C*_*P*_ ( 1−ε)<1 which is always true based on the definitions of a gene-for-gene model (*e.g.* Leonard 1977; Tellier & Brown 2007). The value of the internal equilibrium point defines which allele has a higher frequency in host and parasite populations during the cycles. In the typical GFG parameter space, susceptible hosts and infective parasites have higher frequencies (Leonard 1977, Tellier and Brown 2007).

Only negative indirect frequency-dependence (niFDS) occurs under a geometric seed bank (Tellier & Brown 2007b, 2009), thus the internal equilibrium is unstable (Fig. S3A). This means that allele frequencies are diverging from it, and the coevolutionary cycles progressively increase in period and amplitude (Fig. S3B). As such, the unstable coevolutionary cycles are unpredictable for the plant population at a given generation. The coevolutionary parameters impact the characteristics of the unstable cycles as follows. An increase of the costs *C*_*H*_ (and respectively *C*_*P*_) reduces the advantage of resistance (respectively infectivity), resulting in a shorter period and smaller amplitude of the cycles (see Tellier & Brown 2011, for an approximation of the period of cycles). A higher disease severity *s* also accelerates the cycles (shorter period) but increases their amplitude. Secondly, the seed bank strategy shapes the cycles; a long seed banking persistence *m* increases the period of the cycles, as does an increase of the germination fraction *b*_*0*_ (Fig. S3B).

### Seed banking under GFG coevolution: unlinked loci

Assuming the host locus driving coevolution being independent (genetically unlinked) from the locus determining the germination fraction, means that the susceptible and resistant hosts evolve the same seed banking strategy. In this case, a striking result is the loss of the seed bank when studying the competition of multiple mutants *b*_*0*_* in a resident population *b*_*0*_ (Fig. 1) for moderate disease severity (*s*=0.3 or 0.2) and intermediate or low costs (*C*_*H*_=*C*_*P*_=0.2 or 0.05). This outcome is observed irrespectively of the initial resident strategy and the persistence of the seed bank *m*, with no variance between repetitions (Fig. S4, S5).

**Figure 1:**
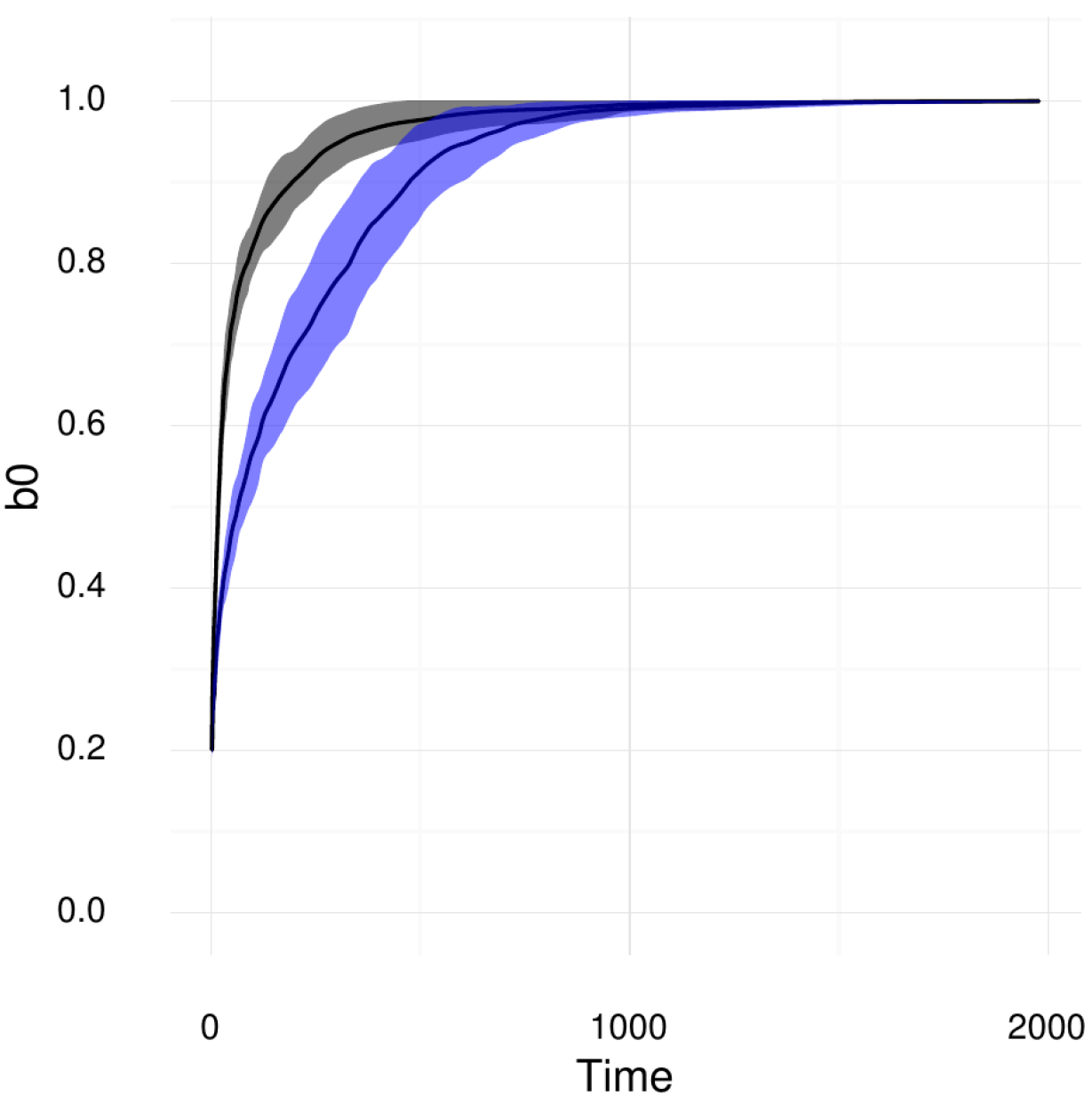
Evolution of the host population mean germination fraction 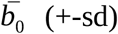 under multiple competition (GFG model, unlinked loci), for short term persistence m=5 (black) and long term persistence m=15 (blue) over 50 repetitions. Costs are fixed to C_H_=C_P_=0.2, s=0.3, and d=0.002. The initial resident value is *b*_*0*_=0.2.

### Seed banking under GFG coevolution: linked loci

Considering non-independent (genetically linked) loci of coevolution and germination and the same coevolutionary costs as the above section, the results are fundamentally different. We observe that resistant and susceptible hosts systematically evolve towards distinct strategies. Surprisingly, a susceptible host does not appear to have any advantage in evolving a seed bank, whereas several germination strategies do evolve for the resistant host. Susceptible hosts always evolve towards a mean population strategy 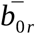 strictly equal to 1 for all persistence values (Fig. 2B-C, Fig. 3B-C), and the pairwise and multiple competition results are fully consistent with each other. We interpret this result as the absence of a seed-bank to be an evolutionary stable strategy for the susceptible host.

**Figure 2:**
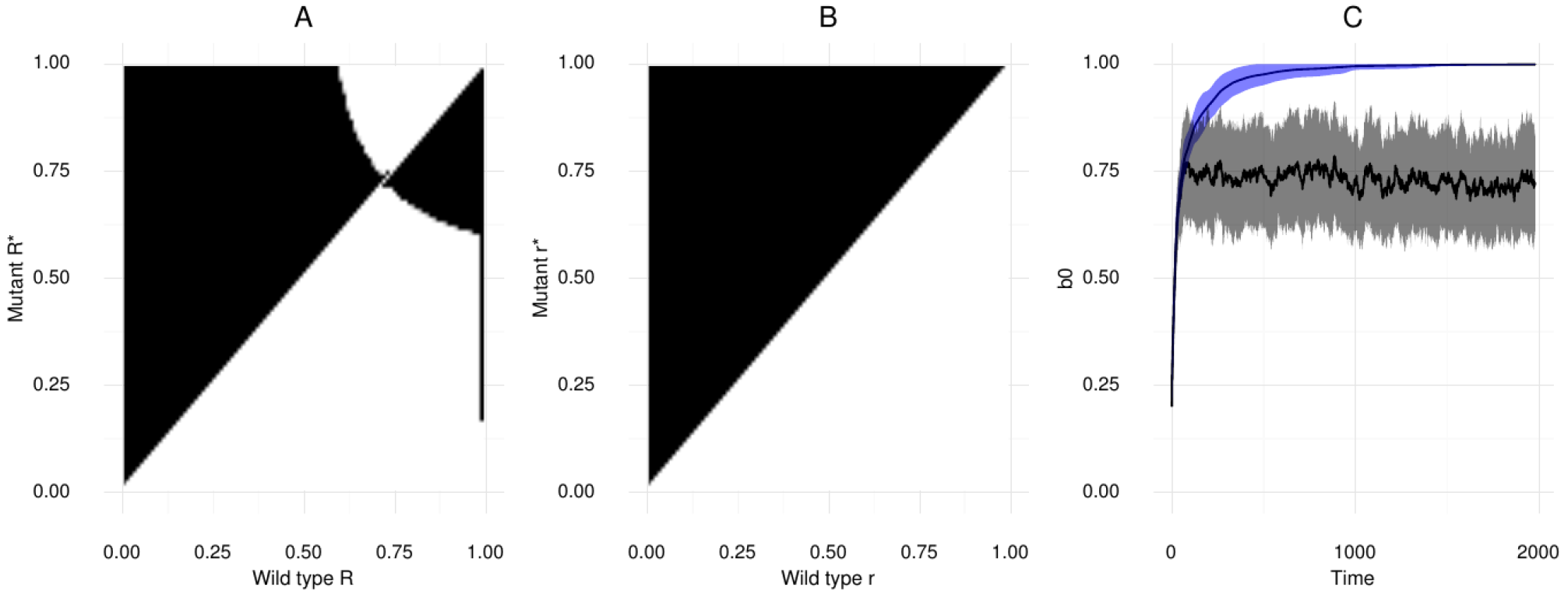
Pairwise invasibility plots (PIP) for GFG model with linked loci, for A) susceptible and B) resistant hosts, with germination fraction *b*_*0*_* of invading phenotypes on the vertical axis, and resident phenotypes b_0_ on the horizontal axis. Hosts have a fixed short term persistent bank m=5, and costs C_H_=C_P_=0.2, s=0.3, and d=0.002. The dynamics are simulated for 1,000,000 generations, the mutant genotype frequency over the last 10,000 generations is superior to 0.9 (black regions), inferior to 0.1 (white regions), or coexists with the resident phenotype (grey regions). C) Evolution of the mean germination fractions (+-sd) of the susceptible 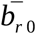 (black) and resistant 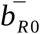 (blue) host under multiple competition with corresponding parameters over 50 repetitions. The initial resident value is b_0_=0.2.

**Figure 3:**
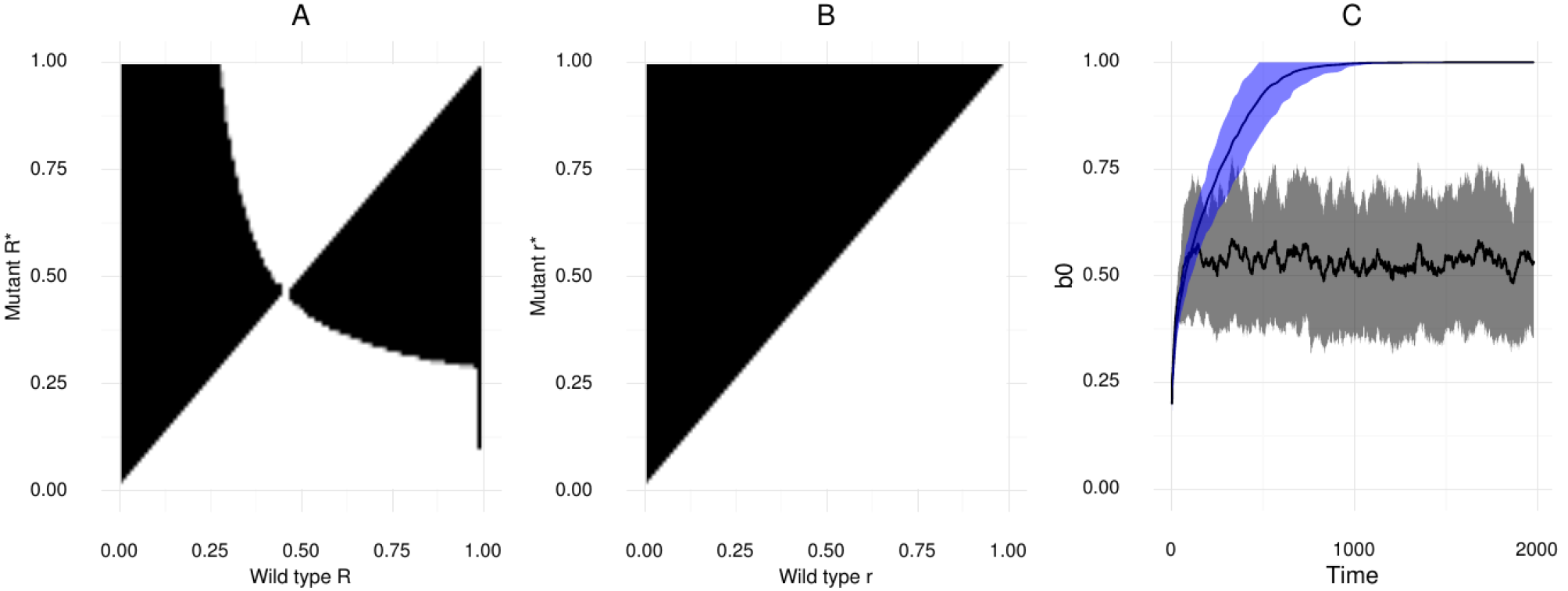
Pairwise invasibility plots (PIP) for the GFG model with linked loci, for A) susceptible and B) resistant hosts, with germination fraction b_0_* of invading phenotypes on the vertical axis, and resident phenotypes b_0_ on the horizontal axis. Hosts have a fixed long term persistent bank m=15, and costs C_H_=C_P_=0.2, s=0.3, and d=0.002. The dynamics are simulated for 1,000,000 generations, the mutant genotype frequency over the last 10,000 generations is superior to 0.9 (black regions), inferior to 0.1 (white regions), or coexists with the resident phenotype (grey regions). C) Evolution of the mean germination fractions (+-sd) of the susceptible 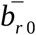 (black) and resistant 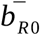 (blue) host under multiple competition with corresponding parameters over 50 repetitions. The initial resident value is b_0_=0.2.

The outcome of seed banking evolution for the resistant host is more complex, and depends on the coevolutionary parameters and on the persistence of the seed bank. For moderate costs (*C*_*H*_=*C*_*P*_=0.2), the result of the pairwise competition shows a range of values, from *b*_*0R*_=0.72 to *b*_*0R*_=0.74, where germination strategies are able to invade and be invaded by one another (Fig. 2A). We interpret this region 0.72< *b*_*0R*_ < 0.74, as consisting of strategies optimising the resistant host’s geometric mean fitness. This supposition is confirmed by simulations under the multiple mutants competition method (Fig. 2C, Fig. S6A-B), in which the resistant host evolves towards germination fractions ranging from *b*_0_≃0.62 to *b*_0_≃0.88, with a mean strategy fluctuating around *b*_*0R*_=0.75 (Fig. 2C). Note that the boundaries of the region are wider than under the pairwise competition prediction (Fig. 2A). We explain it as a consequence of noise inherent to the sampling of mutants in the second method. Although not measured here, as we only account for the resistant mean strategy, the combination of the two approaches suggests that polymorphism could evolve, with the coexistence of close strategies belonging to the optimising region (0.71< *b*_*0R*_ < 0.76). A similar pattern is observed with the increased persistence of the seed bank (*m*=15 in Fig. 3), but the optimum strategy shifts towards the value *b*_*0R*_=0.47 both for the pairwise competition method (Fig. 3A), and for the multiple mutant competitions (Fig. 3C, Fig. S6C-D), albeit being more variable in the latter. The investment in the seed bank is thus stronger with increased persistence. Here, around half of the seeds produced enter the long term bank compared to less than a quarter for a shorter bank (*m*=5).

Considering small costs (Fig. S7-9) the range of germination fractions strategies optimizing the resistant fitness is in line with those observed for moderate costs (Fig. 2-3), though our simulations show a greater variability around the mean resistant strategy in Fig. S7C, S8C, S9 compared to Fig. 2C, 3C, S6. As small costs give a greater advantage to resistant hosts, slower coevolutionary cycles are generated. Consequently, when considering resistant genotypes, the relative advantage of a given strategy against another one is small, explaining the observed variability. Furthermore, for high germination fraction values (>0.79), the frequency of resistant hosts becomes negligible (under 10^−300^). This latter outcome is biologically unrealistic, and we exclude this range of values from our results (Fig. S7-8).

To summarize our results so far, we show the loss of the seed bank at moderate disease severity in the case of unlinked loci, while this loss is observed only for the susceptible host when considering linked loci. This is due to the fitness asymmetry between the two host types that is inherent to the GFG model and acting on two levels. Firstly, the susceptible host is constantly infected by the two types of parasites. As we demonstrated in the SI text S1 model A3 for reduced fitness fluctuations a weak to no investment in the bank is optimal. Secondly, the susceptible host is the most common type over time, whereas the resistant host is only infected by the virulent parasite and undergoes extreme fitness fluctuations – fast increase of fitness but of short period (Fig. S3B). Thus the evolution of the seed banking strategy is governed by the susceptible host under unlinked loci. The fitness asymmetry between resistant and susceptible genotypes depends on the different coevolutionary parameters (costs and disease severity), and is released by switching to a Matching Allele (MA) interaction.

We run simulations of multiple mutant competitions for increasing coevolutionary costs (*C*_*H*_=*C*_*P*_ up to 0.4) together with increasing disease severity *s* (0.1, 0.3, 0.6 and 0.9). Each combination of parameters results in specific coevolutionary cycle characteristics, for instance the cycling periods are short for high costs *C*_*H*_=*C*_*P*_0.4 in comparison with small costs C_H_=C_P_=0.05. In the first case, the fitness asymmetry between host types is reduced. The speed of the coevolutionary cycles has a direct incidence on the evolution of seed banking strategies (Fig 4, S10) as we now observe contexts where both susceptible and resistant hosts evolve a seed banking strategy. This is observed for moderate to strong costs of resistance and infectivity (*C*_*H*_=*C*_*P*_>0.1) and for strong disease severity (*s*=0.6 and/or 0.9). In these contexts, the coevolutionary cycles are fast. Again, the investment in the seed bank is stronger for long term seed banks. The results under unlinked loci now change drastically (Fig 4, S10), as now the host population can evolve seed banking. Indeed the host population strategy is still mostly influenced by the susceptible host type, which can evolve *b*_*0*_ <1.

**Figure 4:**
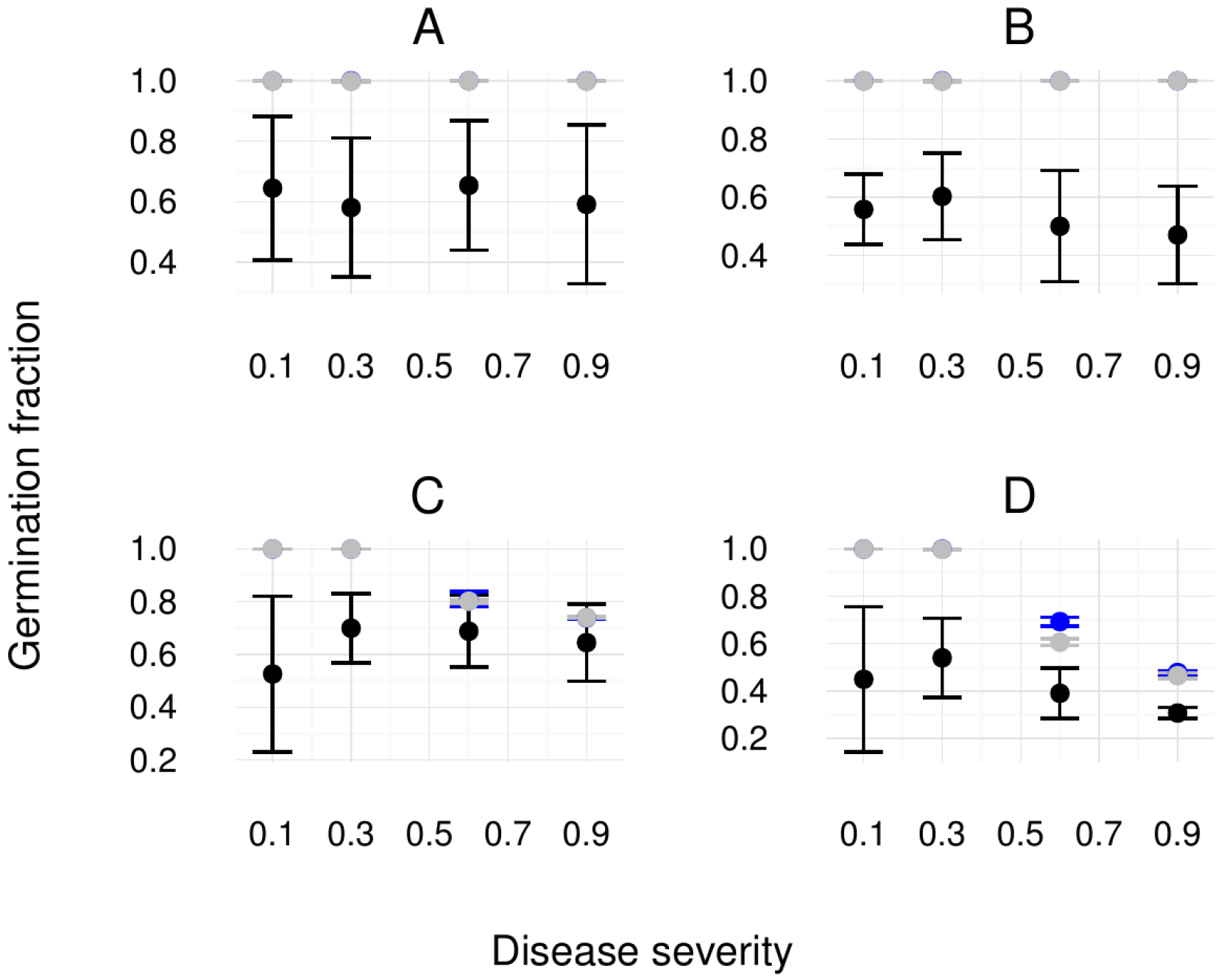
Population mean germination fraction (grey), resistant mean germination fraction *b*_*0R*_ (black) and susceptible mean germination fraction *b*_*0r*_ (blue) as a result of the multiple mutant simulation process (2×10^16^ generations) and function of the disease severity *s* (x-axis). The GFG model has costs of resistance and infectivity fixed to *C*_*H*_=*C*_*P*_=0.05 (A-B) and *C*_*H*_=*C*_*P*_=0.2 (C-D). The seed bank persistence *m* is fixed to 5 years (A-C) and 15 years (B-D). Error bars indicate variation over the 50 repetitions per parameter set. The initial germination fraction is *b*_*0R*_=*b*_*0r*_=0.2.

### Seed banking under Matching allele coevolution

We modify our model to include a Matching Allele recognition matrix (SI; text S3) in which both hosts are infected by their specific parasite and have equivalent fitness fluctuations. We then observe that hosts evolve a seed banking strategy (Fig. S11), and moreover evolve towards the same region of optimal strategies (Fig. S12-14) depending on the persistence of the seed bank. The results are consistent for both types of simulations considered (details in SI). These extended results under the MA model demonstrate that the evolution of seed banking is indeed due to coevolutionary dynamics.

## Discussion

Our results show that seed banking in hosts can evolve in response to parasite pressure and coevolutionary dynamics as a temporal bet-hedging strategy, especially if costs of the resistant and infectivity alleles and disease severity are strong. We have chosen not to separate between the ecological (coevolution) and the evolutionary (evolution of seed banking) time scales in our model, as there is a feedback between the evolution of the germination rate *b*_*0*_ and the coevolutionary dynamics – they do not reach a stable equilibrium or limit cycles stable in period and amplitude. We therefore use simulation methods to demonstrate our hypothesis.

Host genotypes undergo different coevolutionary dynamics depending on their asymmetric or symmetric genetic interaction with parasites (GFG or MA), and as a result genotypes can evolve distinct seed banking strategies. In the GFG model, generalist (*i.e.* susceptible) hosts are found to be the most common type with higher geometric mean frequency than specialist (*i.e.* resistant) hosts. Therefore, under genetic independence of the coevolutionary and germination loci, selection is driven by the generalist host. In addition, the range of seed banking strategies evolving is shaped by the characteristics of the coevolutionary cycles (period and amplitude). Hosts evolve a strict non seed banking strategy (*b*_*0*_=1) for slow coevolutionary cycles due to small costs of resistance and infectivity (C_H_ and C_P_) together with low disease severity (*s*), while various strategies evolve for fast cycles (*b*_*0*_<1) due to moderate to high costs and disease severity. However, under genetic disequilibrium, the resistant host always evolves seed banking while only the susceptible strategy is influenced by the speed of the coevolutionary cycles. Hosts undergoing identical period and amplitude of their coevolutionary dynamics (MA) evolve towards the same seed banking strategies (*b*_*0*_<1).

The fixed seed bank persistence constrains the optimal investment in the bank. Seed banking keeps memory of past selective events, and is most effective when the temporal window covered by the seed bank length (*i.e.* persistence *m*) is large enough with regard to the period of the environmental cycles (Templeton & Levin 1979 and our results in SI text S3). For both GFG and MA models, cycles of host-parasite coevolution show periods much larger than the fixed seed bank persistence *m*. Under our GFG model, the investment in the seed bank is then stronger (*i.e.* higher fraction of seeds entering the bank) for long term persistent banks (*m*=15) than for short term banks (*m*=5); this is particularly visible for moderate to high coevolutionary costs. Decreasing the cost of the specialist genotypes or the disease severity increases the frequency of resistant hosts over time, which in return extends the periods of the coevolutionary cycles and increases the time that allele frequencies remain along the boundaries (defined as frequency of 0 or 1). Although the range of optimal germination fractions is the same as for higher costs, our simulations show hence more variation. As coevolutionary cycles are faster for MA, the investment in the bank is found to be higher (*b*_*0*_=0.6 and *b*_*0*_=0.4 for long and short term persistence, respectively). Host fitnesses being strictly equal with respect to the interaction with parasites, the optimal range of strategy is in this case independent of the genetic linkage. Unstable coevolutionary dynamics can therefore generate bet-hedging.

We make further predictions about the evolution of persistence. If seeds are not constrained physiologically or mechanically, persistence would probably evolve towards extreme values together with low germination fractions to maximise the temporal coverage of the cycling dynamics. An age structure due to perenniality or to a seed bank with different trade-off shapes between the age of a seed and its germination probability, does affect the coevolutionary dynamics (Tellier & Brown 2009) and would influence the evolution of the bet-hedging strategy.

Disease prevalence can also be stochastic and generate equivalent pressures as the abiotic variability usually assumed (Lewontin & Cohen 1969). A strong dependence between the environment, the disease prevalence and disease severity is a common feature to many plant and also invertebrate hosts (the so-called disease triangle in plant disease, Agrios 2005). We thus raise the question whether it is more efficient to evolve resistance to a pathogen or to invest in seed banking. In our infinite size model, assuming a stochastic disease prevalence (but fixed each year) is equivalent to the model of two competing alleles exhibiting stochastic fitness derived in the supplement (SI Text S2). As previous results for abiotic variable environments demonstrated that bet-hedging evolves only when extreme variations in fitness are observed between years (Cohen 1966, text S3), we predict that seed banking would show more benefits than resistance only for extreme stochastic prevalence together with strong disease severity, or assuming high costs of resistance. A second question regards the potential for evolving bet-hedging strategy in parasites under coevolution. Indeed bet-hedging strategies also evolve in parasites, such as low virulence in parasites transmitted by vectors in fluctuating environments (Nguyen *et al.* 2015). Thus we speculate that unstable cycles of coevolution could also generate bet-hedging in parasites, promoting the existence of dormant survival strategies within or outside hosts.

We finally derive the following predictions to test our model. Firstly, an experimental coevolution set up with phage and bacteria, could investigate whether bacteria under coevolutionary pressures evolve a bet-hedging strategy for dormancy depending on their resistance genotype (*e.g.* combining approaches by (Poullain *et al.* 2008; Beaumont *et al.* 2009; Betts *et al.* 2014). Secondly, species in disturbed habitats forming transient populations (*e.g.* metapopulations), may evolve seed banks as bet-hedging strategy for persistence and/also in trade-offs with dispersal (Vitalis *et al.* 2013). Such species may not be in tight coevolution with pathogens, due to their inherent transient nature. However, candidate species to evolve bet-hedging in response to coevolution would be found in more stable habitats (temperate climate) which are pervasive for infections (Jousimo *et al.* 2014). In addition, for seed banking species showing more stable populations in time and a spatial structure, we predict that populations under higher pathogen pressure should exhibit higher resistance (Soubeyrand *et al.* 2009; Jousimo *et al.* 2014) alongside with longer seed banking persistence or lower germination fraction.

A key step to further test our predictions is to disentangle the effect of different selective pressures in driving the evolution of bet-hedging in space (dispersal) or in time (seed banking), even though these belong to a continuum of strategies but rely on different physiological adaptations (Buoro & Carlson 2014). Obviously, several selective pressures due to variation of biotic and abiotic factors generate the condition for bet-hedging to occur and are acting concomitantly on species, perhaps even with fluctuating strength or importance over time and space. Coevolution, as we described, is a slow mechanism, probably acting at longer evolutionary scales (over several cycles) than drastic environmental stochasticity, but we expect that combinations of several variable abiotic and biotic factors would accelerate the evolution of bet-hedging.

The choice for a modelling approach assuming fixed population size is dictated by the core assumption of bet-hedging theory: the environmental unpredictability. Our model ensures that coevolutionnary dynamics are strictly driven by niFDS, leading to unstable cycles, which constitute an unpredictable environment for hosts. However, our results may not apply to all host-parasite systems because our current framework dismisses epidemiological and population dynamics and the potential for eco-evolutionary feedbacks. The latter are known to generate direct frequency-dependent selection, resulting in coevolutionary dynamics exhibiting stable limit cycles or damping off towards a stable polymorphic internal equilibrium point (Tellier & Brown 2009; Ashby & Boots 2017). As such, an epidemiological approach cannot generate the evolution of bet-hedging in the strict sense (Boer 1968), and is outside the scope of this study. Epidemiological dynamics produce strong eco-evolutionary feedbacks when parasites survive only within their hosts and either kill them to be transmitted or induce large fitness damage to their host (e.g. Gokhale *et al.* 2013, Ashby & Boots 2017). Considering the diversity of parasite life cycles and life history traits as well as the influence of the abiotic environment on host-parasite interactions, it is conceivable that strong eco-evolutionary feedbacks may not always apply. As a consequence, short term epidemiological models may not accurately predict the longer term coevolutionary dynamics and cycles over thousands of generations, the time scale at which seed banking strategies may evolve. The simple model based on fixed population sizes and longer time scale we use here represents a first necessary step in order to disentangle the effect of coevolutionary dynamics *sensu stricto* (*i.e.* changes in allele frequencies in interacting species) from that of eco-evolutionary feedbacks on the evolution of seed banking. Further studies should include the effect of population dynamics and epidemiology in our framework, to investigate not only if seed banking evolves under such conditions, but also the interaction between epidemiological dynamics and the buffering effect of the seed bank on population sizes.

## Acknowledgements

MV and AT acknowledge funding from DFG Grant TE809/1-1. We thank Daniel Zivkovic for help with Mathematica computations, and Frederic Hamelin for discussions and detailed comments on the manuscript.

